# Reconstructing the complex evolutionary history of the fused Aco2 gene in fission yeast

**DOI:** 10.1101/2019.12.16.879015

**Authors:** Steven Sun, Scott William Roy, Noelle Anderson

## Abstract

The Aco2 gene of *Schizosaccharomyces pombe* was formed by gene fusion between curious partners, namely genes encoding the enzyme aconitase and a mitochondrial ribosomal protein. In addition to a full-length transcript, a truncated mRNA encoding only the N-terminal aconitase domain is produced by polyadenylation at an internal site. Protein products of the gene are found in the nucleus, mitochondria and cytoplasm, consistent with the presence of multiple subcellular targeting signals. To reconstruct the evolution of this complex gene, we studied homologous genes from a range of related species. We find evidence for a dynamic history within Taphrinomycotina, including: early evolution of a nuclear localization signal; creation of a 3’ intron that could be involved in regulating subcellular targeting; evolution of multiple peroxisomal targeting signals in different lineages; and recurrent gene loss. We present a likely stepwise model for the evolution of this remarkable gene and discuss alternative scenarios.

---

A central goal of biology is to understand the relationship between structure and function. Structure-function relationships concerning eukaryotic nuclear genomes remain obscure, as the genomes are littered with noncoding sequences and often undergo alternative processing by alternative splicing, polyadenylation and RNA editing (Albertin et al. 2015, Raj and Blencowe 2015, Taliaferro et al. 2016). The incidence of alternative transcript processing that has functional significance remains unclear. Within animals and plants, well studied cases show clear functions for alternative processing, however the functional significance of the large majority of uncharacterized events remains debated (Albertin et al. 2015, Irimia and Blencowe 2012, Landry et al. 2003, Di Giammartino et al. 2011, Tress et al. 2017, Blencowe 2017, Syed et al. 2012). In other eukaryotes, incidences of alternative processing are often lower and no evidence for function exists for the overwhelming majority of cases (McGuire et al. 2008, Hon et al. 2013, although see e.g., Chen et al. 2014). Within the entirety of fungi, only a small number of alternative processing events have any evidence for function, most of which involve alternative subcellular targeting (Jung et al. 2015, Freitag et al. 2012, Steibler et al. 2015).

One of the most striking known cases of alternative processing in fungi involves the Aco2 gene of *S. pombe*. This gene was formed by fusion between genes encoding two functionally very different proteins: (i) aconitase, the enzyme that catalyses the isomerization of citrate to isocitrate in the tricarboxylic acid cycle; and (ii) L21, a core mitochondrial ribosomal protein. Interestingly, in addition to a full-length form, the fused gene expresses a truncated form including only the 5’ aconitase-encoding region, which is produced by alternative use of an internal polyadenylation site (see Figure 1a; Jung et al. 2015), leading to production of a (non-genomically encoded) stop codon.

**FIgure 1.**
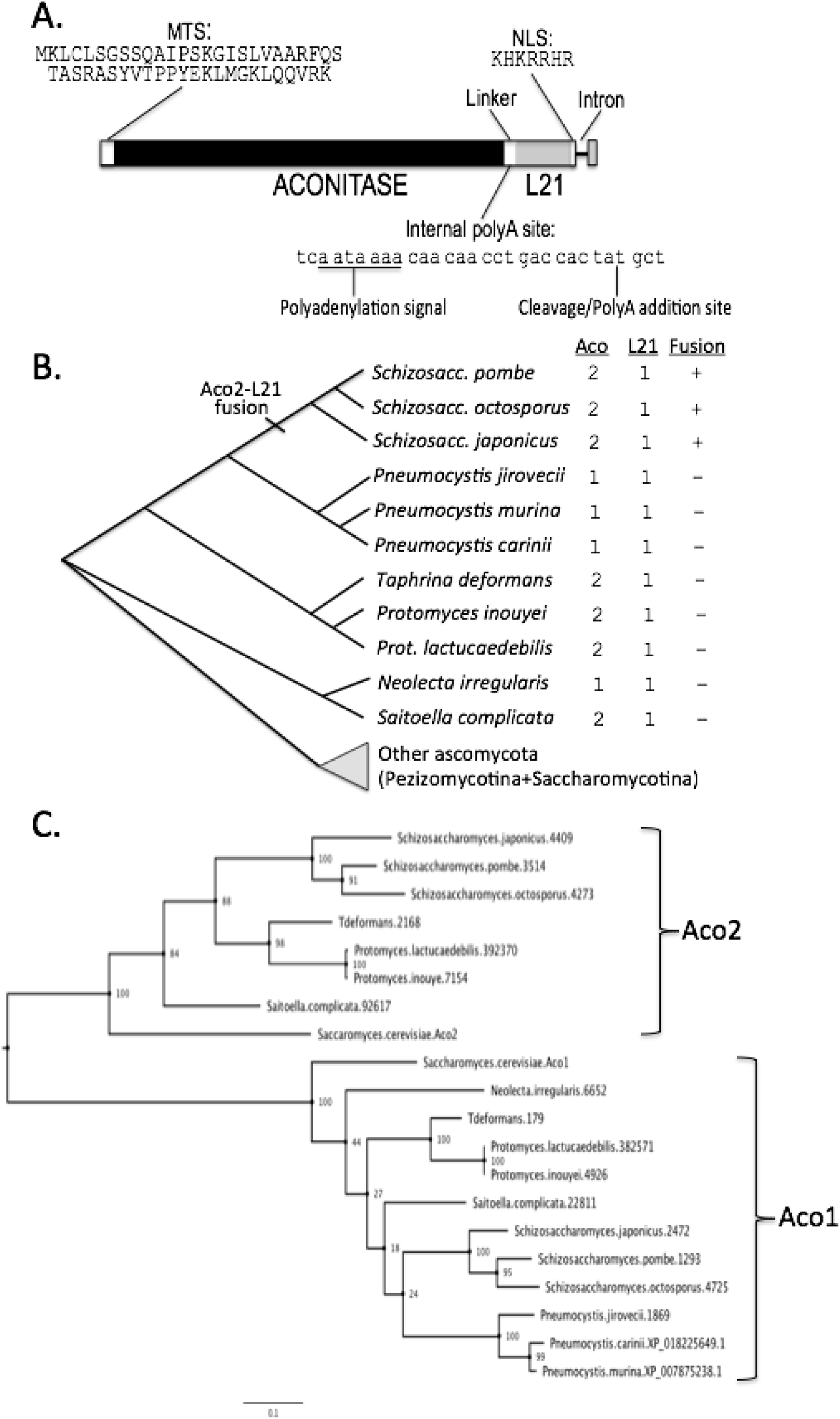
Previous results and gene features for 11 species of Taphrinomycotina fungi. A. Summary of previous results from Jung et al. The S. pombe Aco2 gene contains fused Aconitase and L21 domains, as well as an intervening linker region. Additional features include an N-terminal mitochondrial targeting peptide (MTS), an internal alternative polyadenylation site that converts a genomically encoded UAU codon into a UAA stop codon, a nuclear localization signal and a single 3’ intron. B. Number of aconitase genes, L21 genes and presence/absence of a fused aconitase-L21 gene for 11 Taphrinomycotina species with available genomes. C. Gene tree for 18 aconitase genes in the 11 Taphrinomycotina species, as well as Aco1 and Aco2 genes for *S. cerevisiae*, showing clear Aco1 and Aco2 clades. Bootstrap support values are given.

While fusion of genes with such different core functions seems peculiar, the finding becomes less surprising in light of evidence for direct interaction between the non-fused Aco2 gene of *S. cerevisiae* and the mitochondrial ribosome. Also of likely importance are reports of diverse secondary (“moonlighting”) functions for aconitase enzymes (e.g., Klausner and Rouault 1993). For instance, under low iron conditions, an aconitase protein in animals undergoes a conformational change, producing a non-catalytic form that regulates genes involved in iron metabolism by binding their mRNAs (Basilion et al. 1994). Another example involves ACO1 in *S. cerevisiae* (Chen et al. 2007), which can bind and stabilize mitochondrial DNA upon disassociation of its 4Fe-4S iron-sulfur (Chen et al. 2007).

Here we reconstruct the evolutionary history of the fused Aco2-L21 gene and its homologs.

## Summary of previous results

Figure 1a summarizes results from previous studies. Jung et al. (2015) performed a detailed study of the Aco2-L21 fusion gene, highlighting: (i) existence of multiple subcellular targeting signals including an N-terminal mitochondrial targeting sequence (MTS) and a C-terminal nuclear localization signal (NLS); (ii) an internal alternative polyadenylation site leading to production of a polyadenylated mRNA encoding only the aconitase domains; and (iii) targeting of the different isoforms to various subcellular locations.

## Fusion and losses of Aco2

We downloaded the genomes and proteomes for 11 Taphrinomycotina species from JGI (Figure 1b) and performed BLASTp and tBLASTn searches against all 11 species to determine gene presence and fusion status for each species. For each species, a single sequence homologous to L21 and either one or two aconitase homologs were found (Figure 1b). As previously reported, the Aco2-L21 fusion was restricted to *Schizosaccharomyces* (Figure 1b). Phylogenetic reconstruction of identified aconitase protein sequences yielded two strongly-supported clades, one corresponding to Aco1 and the other to fused and unfused Aco2 proteins (Figure 1c). The Aco2 clade lacked representatives from the three studied *Pneumocystis* species and *Neolecta irregularis*, suggesting multiple losses. The presence of an unfused L21 copy in *Pneumocystis* suggests the fusion occurred after the *Schizosaccharomyces-Pneumocystis* divergence, however earlier fusion followed by partial gene degradation in *Pneumocystis* cannot be excluded.

## Turnover of subcellular targeting

Scrutiny of Taphrinomycotina Aco2 genes revealed a conserved classical C-terminal peroxisomal targeting signal (PTS1) in a clade of three species (Figure 2a). This is curious, since proteome-level studies have not found aconitase in peroxisomes (Schäfer et al. 2001, Managadze et al. 2010), and a search for classical PTS1 tripeptides ([A/S]-[R/K]-[L/I]) revealed that such signals are rare across Ascomycota aconitase genes (data not shown). Interestingly, the fused Aco2-L21 genes contain a conserved candidate N-terminal peroxisomal targeting signal (PTS2), with close match to the reported PTS2 consensus sequence [R/K]-[L/V/I/Q]-X-X-[L/V/I/H/Q/T]-[L/S/G/A/K/T]-X-[H/Q]-[L/A/F/I/V] (Petriv et al. 2004) (Figure 2b). Nuclear localization signals (NLS) searches using NLS mapper (Kosugi et al. 2009) revealed strong candidates in nearly all L21 genes, suggesting an ancestral NLS (bold sequences in top of Figure 2c).

**Figure 2.**
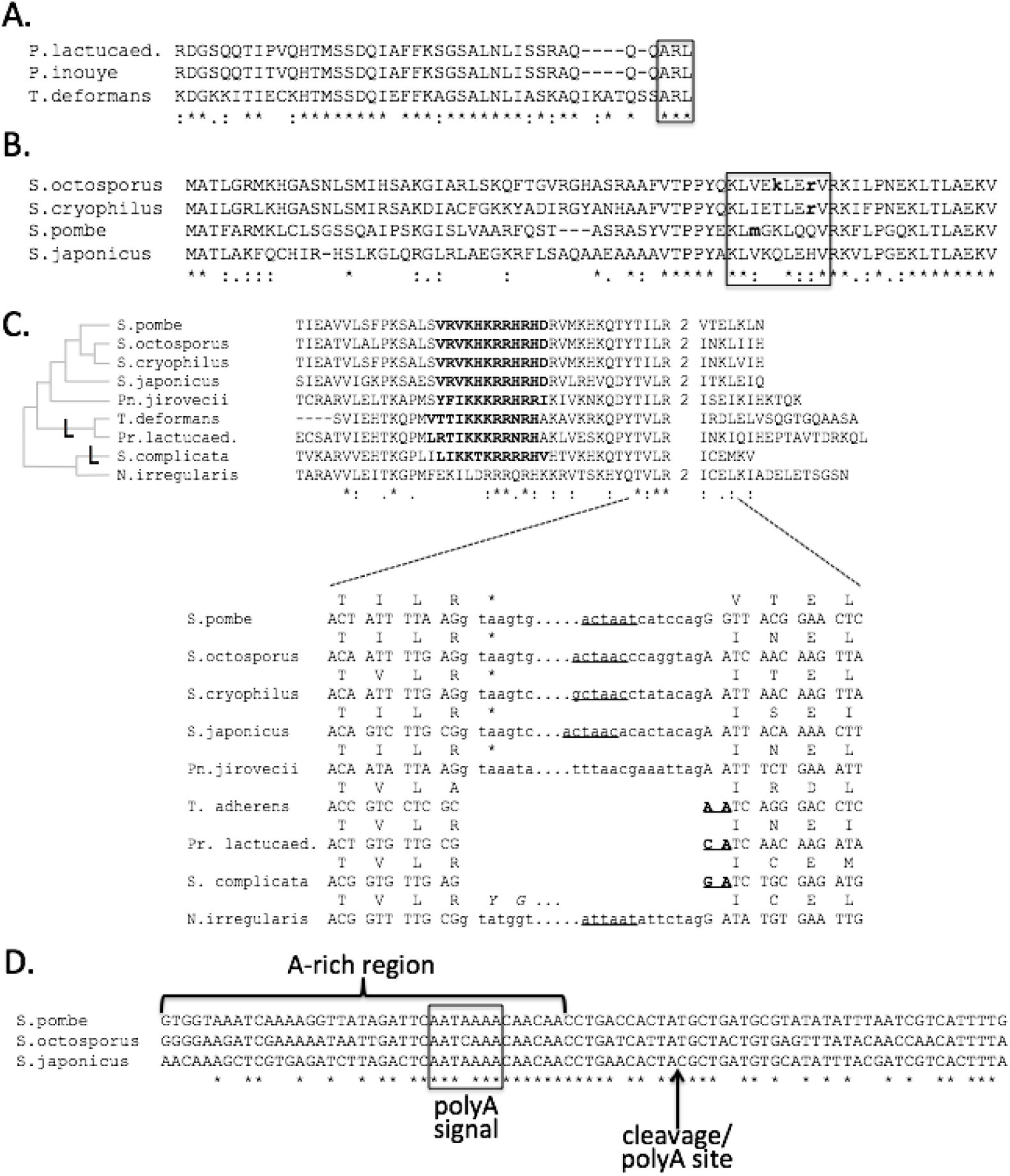
Subcellular localization signals and transcript processing of fused and unfused Aco2 and L21 genes in Taphrinomycotina. A. Putative C-terminal peroxisomal targeting signals (PTS1) in unfused Aco2 genes of three Taphrinomycotina species. B. Putative peroxisomal targeting signals proximal to N terminus (PTS2) in fused Aco2-L21 gene of *Schizosaccharomyces* species. C. C-terminal sequence and gene structure of fused and unfused L21 genes, showing NLS (bold) and presence/absence of a phase 2 intron near the stop codon (“2”), with multiple inferred intron losses (“L”) shown on the phylogeny. Nucleotide sequences for corresponding region shown below, highlighting intron boundaries and putative branchpoint sequence (underline); in-frame stop codons for *Schizosaccharomyces+Pneumocystis* (“*”); and lack of G[T/C] splice boundaries for other species (bold underline). D. Conservation of polyA signal and cleavage/polyA sites for alternative internal polyadenylation site. The region with elevated A content which could represent a degraded poly-A tale from a retroposition event is noted.

## A 3’ intron in L21

We noted an intron near the stop codon in both fused *Schizosaccharomyces* genes and unfused L21 genes of a subset of Taphrinomycotina species (Figure 2c). For *Schizosaccharomyces’* sister genus (*Pneumocystis*), no 3’ intron was annotated. However, we noted candidate splicing signals (Figure 2c). Mapping of available RNA-seq Illumina data confirmed splicing, with 17 spliced reads and zero unspliced. Given Aco2-L21’s variety of subcellular targeting signals, we speculated that alternative splicing of the 3’ region could produce differentially-targeted alternative protein isoforms, however, available RNA-seq reads from *S. pombe* revealed efficient splicing under a variety of conditions (≥98% of observed reads spliced).

## Fusion of Aco2 and L21

To probe the mechanism of gene fusion, we studied intron-exon structures. Like many Ascomycetes species, in *Protomyces lactucaedebilis*, the predicted L21 gene does not contain introns. While tBLASTn searches against the genome indicate the presence of an L21 gene in both *P. inouyei* and *T. deformans*, these genes are not annotated and thus their intron-exon structure remains unknown. By contrast, L21 genes in *Pneumocystis* species contain an intron ∼25 codons downstream of the start codon, however this region is not well conserved across species and so could have arisen by any number of mechanisms. This region of the L21 protein is not well conserved across these species, nor across fungi in general, making it impossible to characterize the sequence or structure of the L21 gene involved in the fusion. Similarly, unfused Aco2 genes do not yield much insight into the structure of the Aco2 gene involved in the fusion. The AKL putative PTS1 signal tripeptide found in the *Taphrina/Protomyces* clade is not present at the homologous position of other species, however, it is not possible to tell whether this reflects the sequence at the time of divergence, subsequent sequence evolution, or a more complex event involving fusion of only a partial Aco2 gene to L21.

Scrutiny of the fused and unfused genes also did not reveal clear evidence as to whether the fusion might have occurred by retrotransposition. Two signatures of retrotransposition are loss of introns and the presence of a polyA tail. In general, unfused Taphrinomycotina Aco2 genes are intronless, thus the absence of introns within the Aco2 section of Aco2-L21 is not informative as to whether the Aco2 gene might have retroposed into proximity with L21. As for the presence of a polyA tail, a fairly A-rich sequence overlapping the internal alternative polyadenylation site (Figure 2d) is found downstream from the end of the conserved Aco2 sequence, as would be expected by retroposition. However, the sequence is not overwhelmingly A-dominated (49-51% of nucleotides are A’s across species), and such short A-rich sequences are not uncommon (e.g., present in ∼30% of *S. pombe* genes), thus this is also not strong evidence for retroposition.

In short, all that can be said about the fusion event is that in extant fused copies, conserved N-terminal Aco2 sequence is linked to conserved C-terminal L21 sequence by a stretch of sequence that does not show recognizable homology to any sequences in non-fused L21 or Aco2 sequences and is faster evolving than surrounding the L21 and Aco2 domains (Figure 2d).

## Evolution of polyadenylation sites

Finally, we studied the evolution of the region surrounding the polyadenylation site. Comparing within *Schizosaccharomyces*, we found conservation across the genus of the consensus polyadenylation site AATAAA, as well as the TA[C/T] codon that acts as the cleavage/polyA-addition site (Figure 2d). Comparison with non-*Schizosaccharomyces* species showed that the *S. pombe* polyadenylation motif and cleavage site fall within non-conserved regions (sequence putatively associated with the gene fusion). Scrutiny of the genomic regions downstream of the stop codon for species in the *Taphrina*/*Protomyces* clade failed to reveal potential polyadenylation motifs (data not shown).

## Reconstruction of the evolutionary history of Aco2-L21s

These results allow us to reconstruct the stepwise evolution of the fused *Aco2-L21* gene (Figure 3). First, early in the evolutionary history of Taphrinomycotina, L21 acquired a nuclear localization signal. We speculate that nuclear targeting of this protein was associated with a newly-evolved moonlighting function, perhaps involving redeployment of a RNA-binding function initially evolved in mitochondria. Possibly second was the evolution of peroxisomal targeting of Aco2, though timing remains uncertain since it is unknown whether the PTS1 signal was ancestral and replaced by PTS2 or vice versa, or whether the signals arose independently. Finally, multiple changes occurred in the lineage leading to the ancestor of *Schizosaccharomyces*, including: (i) gene fusion by an unknown mechanism to give rise to a gene structure with the single 3’ intron from the ancestral L21 gene; (ii) evolution of an internal polyadenylation site; and (iii) evolution of a 3’ intron.

**Figure 3.**
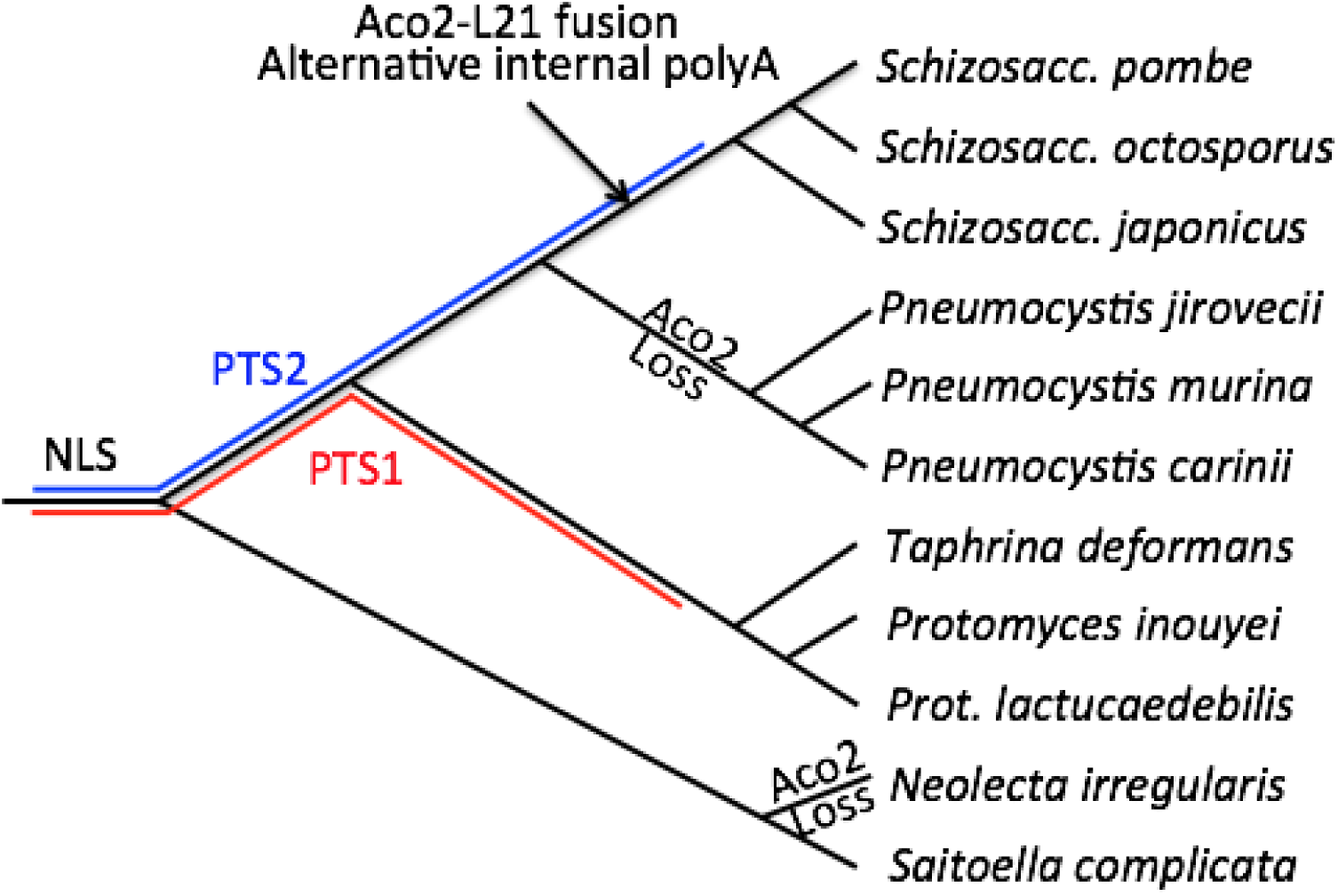
Evolutionary history of Aco2 and L21 genes within Taphrinomycotina. NLS/PTS1/PTS2 represent the timing of origin of cellular localization signals, with ambiguities of timing of PTS origin represented by colored bars. Multiple losses of Aco2 genes are indicated, as are transformations inferred to have occurred in the *Schizosaccharomyces* ancestor.

## The complex evolutionary history of Aco2

It is of note that Aco2 of ascomycetes fungi appear to have a complex evolutionary history. In *S. cerevisiae*, Aco2 is implicated in lysine biosynthesis, performing a reaction associated with homoaconitase in other organisms, while Aco1 performs other various functions, notably in the tricarboxylic acid (TCA) cycle and glyoxylic acid cycle (GAC) (Regev-Rudzki et al. 2005). In contrast, Aco2 in filamentous fungi is not required for lysine biosynthesis, nor for any of the functions performed by Aco1 in *S. cerevisiae* (Fazius et al. 2012). Thus, the subcellular targeting and fusion history of Aco2 occurs in the context of a gene that appears to show peculiar flexibility (particularly for a core enzyme) within ascomycetes fungi more generally. For instance, peroxisomal targeting of Aco2 in Taphrinomycotina is surprising given that Aco1 is involved in the GAC of *S. cerevisiae*, but takeover by Aco2 of functions performed by other homologs is precedented given its role in biosynthesis in *S. cerevisiae*.

## Turnover of peroxisomal metabolism and parallel evolution of subcellular targeting

Evidence for peroxisomal targeting of Taphrinomycotina aconitases is somewhat curious. While aconitase is involved in GAC, which takes place in modified peroxisomes of plants and fungi called glyoxysomes, peroxisomal organelles are rich in reactive oxygen species that render aconitase inactive by oxidizing its iron-sulfur clusters (Klausner and Rouault 1993). Not surprisingly, then, aconitase is generally found in the cytosol (as can be other GAC enzymes such as ICL1; Kunze et al. 2006), and metabolic intermediates diffuse across the peroxisomal membrane.

Possibly, PTS signals allow ACO2 to associate with the exterior of the peroxisome, perhaps through tethering to the peroxisomal membrane, however, PTS signals are not evident in previous studies of peroxisomal membrane proteins (Schäfer et al. 2001). Such ‘partial’ targeting could be aided by the presence of weak PTS signals or by competing subcellular localization signals, as seen in various dual-targeted fungal proteins (Ast et al. 2013).

Another intriguing possibility is raised by Haimovich et al. (2016), who proposes co-translational import of peroxisomal proteins occurs via the PTS2 import pathway. If so, regulatory associations of ACO2 with mRNAs of metabolic enzymes (Klausner and Rouault 1993) could have been co-opted for mRNA transport to the peroxisome to aid in co-translational import of the encoded proteins.

## Concluding remarks

These data shed light on the stepwise evolutionary origins of one of the most complex known fungal genes. As genome-wide data for diverse eukaryotes accumulate, this case study will hopefully allow for the identification of recurrent pathways of genic innovation in multifunctional genes and the ways in which gene fusion can serve as a stabilizer of genotypic innovations (as in associating moonlighting genes with their new partners) or as a means of genotypic innovation. Further experimental exploration of the Aco2 gene should help to further illuminate this fascinating case.s

## Materials and Methods

Aconitase homologs from across Ascomycotina were identified by (i) downloading proteome sequences from 243 species from JGI (jgi.doe.gov) and performing local BLASTp searches with e-value cutoff of 10^−20^; (ii) downloading genomic sequences from Taphrinomycotina species from JGI and performing local tBLASTn searches with e-value cutoff of 10^−10^. Sequences were aligned using MUSCLE v3.8.1551 (Edgar 2004) and trimmed using trimAl v1.4.rev15 (Capella-Gutiérrez et al. 2009) in ‘strict’ mode, which automatically sets a trimming threshold based on the similarity score distribution from the alignment. RAxML version 8.2.11 (Stamatakis 2014) was used to reconstruct the phylogeny, using the GAMMA model of rate heterogeneity and a ML estimate of alpha-parameter with 10,000 rapid bootstrap replicates.

All available RNA-seq reads for *P. jiroveci* were downloaded from SRA (BioProject PRJEB3063, ERP001479). All *S. pombe* RNA-seq reads from Saint et al. (https://www.biorxiv.org/content/biorxiv/early/2018/04/23/306795.full.pdf) were downloaded from SRA (PRJEB29696). For each species, all RNA-seq reads were mapped against spliced and unspliced forms of the corresponding Aco2 gene and uniquely mapped reads tallied. NLS searches were performed using the online NLS mapper tool with default parameters (http://nls-mapper.iab.keio.ac.jp/cgi-bin/NLS_Mapper_form.cgi).

